# The role of STN beta oscillations on lower extremity muscle activity in Parkinsonian stepping

**DOI:** 10.1101/2025.01.01.631000

**Authors:** Thomas G. Simpson, Shenghong He, Alek Pogosyan, Fernando Rodriguez Plazas, Michael G. Hart, Rahul Shah, Harutomo Hasegawa, Christoph Wiest, Laura Wehmeyer, Sahar Yassine, Xuanjun Guo, Anca Merla, Andrea Perera, Ahmed Raslan, Andrew O’Keeffe, Marie-Laure Welter, Keyoumars Ashkan, Francesca Morgante, Pablo Andrade, Veerle Visser-Vandewalle, Erlick A. Pereira, Huiling Tan

## Abstract

Freezing of gait (FOG) is a devastating symptom of Parkinson’s disease (PD) often resulting in disabling falls and loss of independence. It affects half of patients, yet current therapeutic strategies are insufficient, and the underlying neural mechanisms remain poorly understood. This study investigated beta oscillation dynamics in the STN during different locomotor states, while examining the effects of levodopa. In particular, it aimed to identify pathological activity by analysing the relationship between the STN and lower limb muscles during stepping. Local field potentials (LFP) in the STN and muscle activity (EMG) of the gastrocnemius and peroneus longus were recorded in 14 PD patients during standing and stepping, ON and OFF levodopa. Levodopa reduced stepping variability, implying improved stepping abilities. Distinct STN beta patterns were observed between stepping and standing, with lower high-beta and higher low-beta during stepping compared to standing, suggesting a distinct role of these frequency bands in motor control during postural and movement states. Levodopa reduced low-beta but increased high-beta activity, highlighting a potential physiological function of high-beta in the STN during standing and stepping. In addition, step-phase specific effects of levodopa included reduced broad-beta band activity in the STN and lower limb muscles during the late-stance and pushing-off phase of the contralateral leg when ON medication. Further analyses suggest that pathological STN activity amplifies muscle activation around movement initiation, potentially reducing the ability of the patient to move freely. These findings offer insight for developing phase-specific stimulation strategies targeting STN beta oscillations during gait.

## 1. Introduction

Freezing of gait (FOG) is a severely debilitating symptom of Parkinson’s disease (PD) affecting around 70% of patients when disease duration exceeds 10 years (Ge et al., 2020). This symptom often leads to a significant loss of independence, a reduction in quality of life, and an increased risk of disabling falls. Current interventions for parkinsonian gait generally offer limited efficacy (Müller et al., 2019; Smith et al., 2021), highlighting the demand for optimised therapeutic approaches. To date, the pathophysiology of FOG is not fully understood as gait networks are highly complex and challenging to investigate. However, research has identified several contributing factors, including altered timing of the gait cycle, insufficient forward progression, and deceleration, which can accumulate and ultimately result in breakdown of the movement (Nieuwboer et al., 2004). Freezing episodes can also emanate from an inability to adapt motor programs, evidenced by challenges with specific gait scenarios, such as turning (Spildooren et al., 2019), movement initiation (Rosin et al., 1997), and movement cessation (Cameron et al., 2010). Structural and functional anomalies in numerous brain areas have also been implicated in FOG, including the frontal cortex, basal ganglia, cerebellum and mesencephalic locomotor region (MLR) (Fasano et al., 2015; Lench et al., 2020; Strelow et al., 2022). However, the underlying neural mechanisms of gait and FOG remain unclear, emphasising the need for further research.

The subthalamic nucleus (STN) has a key role in gait, as it receives ‘hyperdirect’ inputs from motor areas and serves as an important moderator of basal ganglia output (Hamani et al., 2004). It is thought to regulate the integration of cortical and cerebellar information by activating or inhibiting the MLR (Arnulfo et al., 2018; Takakusaki, 2008), which has an active role in initiating and modulating spinal neural circuitry for motor control (Takakusaki, 2017). As the primary surgical target for deep brain stimulation (DBS) in PD, the STN has been extensively studied via electrophysiological recordings and consistently shown to exhibit increased oscillatory activity in the beta (13-30Hz) frequency band, which has been linked to symptoms of rigidity and bradykinesia (Brown, 2007; Little & Brown, 2014). This overactivity can lead to excessive inhibition of the MLR, disrupting the normal initiation and regulation of gait, and potentially contributing to symptoms such as FOG episodes (Mirelman et al., 2019). In addition, prolonged beta burst durations in the STN have been shown to differentiate freezers from non-freezers, with shorter bursts observed in non-freezers (Anidi et al., 2018). Further examination of STN beta modulation within the gait cycle has revealed step-phase specific modulations with fluctuations in STN beta oscillations during stepping possibly reflecting variations in motor output within the gait cycle (Fischer et al., 2018).

Dopaminergic treatment with levodopa has been consistently shown to decrease beta activity in subcortical structures, including the STN, while positively impacting objective measurements of gait such as velocity, stride length, rigidity, and movement initiation (L. Gao et al., 2017; Lubik et al., 2006; Prochazka et al., 1997). However, how levodopa affects the gait-phase related beta-band modulation in the STN remains unclear. Similarly, DBS of the STN, which is thought to attenuate beta activity, has demonstrated efficacy in improving PD motor symptoms, including gait and FOG (Karachi et al., 2019). Recent advancements in adaptive DBS based on beta band activity have further shown promise in ameliorating gait symptoms (Isaias et al., 2024; Wilkins et al., 2024). Nevertheless, the impact of DBS on gait in PD is complex, with most studies reporting improvements but occasional cases observing worsened or newly induced FOG (Gao et al., 2020). These prior findings underscore the critical role of STN beta oscillations in gait regulation and FOG control, emphasising the need for a deeper understanding of these mechanisms to optimise therapeutic strategies.

Apart from abnormal neural activities, pathological muscle activation patterns in the lower limbs have also been linked to FOG (Caliandro et al., 2011; Cantú et al., 2019; Nieuwboer et al., 2004). For example, FOG is associated with a reduction in total EMG activity in lower limb muscles, including the tibialis anterior (TA) and gastrocnemius (GA), as well as shorter durations of muscle activation (Caliandro et al., 2011; Cantú et al., 2019; Nieuwboer et al., 2004). This reduction was accompanied by increased amplitudes of EMG bursts in the TA, suggesting a compensation strategy of pulling the leg into swing. In contrast, more recent research findings indicate that PD freezers exhibit increased muscle activity within the alpha and low-beta bands in both the TA and gastrocnemius GA muscles compared to healthy control during walking (Breu et al., 2022). In addition, beta bursts in the cortical motor network have been found to be associated with increased beta band activities in upper limb muscle activity and cross-muscle phase synchrony in healthy motor control (Echeverria-Altuna et al., 2022;

Simpson et al., 2024). This suggests that motor control involves a coordinated relationship between brain and muscle activity, where alteration of such synchronization may contribute to pathological states in PD. In fact, increases in pathological beta and theta rhythms in the STN have been observed to be followed by a temporal chain of abnormal lower limb muscle firing, detected by EMG (Georgiades et al., 2019). Nevertheless, the precise mechanisms by which abnormal brain activity translates into pathological muscle activation, and how this contributes to gait impairments like FOG in PD, is poorly understood.

This study aims to elucidate the effects of dopaminergic treatment (levodopa) on beta band activity in the STN during stepping in PD. The first objective was to examine how levodopa influences low- and high-beta activity in the STN during standing and stepping, providing insights into the distinct roles of beta oscillations in these locomotion states. Furthermore, this work intends to better describe how STN beta activity and lower extremity muscle activity dynamically change throughout the step cycle and how these changes are modulated by levodopa at different stepping phases. In addition, the relationship between pathological STN beta bursts and aberrant muscle activation patterns during stepping was explored. The ultimate objective is to propose a step-phase-specific stimulation strategy to directly target stepping and gait deficits in PD. To achieve this, 14 PD patients with externalised DBS electrodes implanted in the STN were recruited. Participants completed a stepping paradigm in both OFF and ON levodopa states, while STN local field potentials (LFP), lower extremity electromyography (EMG), and stepping force were recorded.

## 2. Methods

### 2.1 Ethical approval

This experiment was approved by the South Central - Oxford C Research Ethics Committee.

### 2.2 Participants

A total of 14 participants were recruited for the study (*age = 64.5 ± 5.1 (mean ± s.d.) years, disease duration = 11 ± 6.5 (mean ± s.d.) years*), all of whom gave informed consent prior to the experiment. Participants were able to understand and complete the task, although in some cases symptoms were so debilitating, especially when OFF medication, that the paradigm could not be executed in full. In these instances, participants performed for the maximum duration possible. Recordings were conducted 4 to 7 days after the first surgery, with all patients receiving bilateral STN implants (see **Section 2.4**).

### 2.3 Experimental setup

The paradigm consisted of 3 separate segments: rest, standing, and stepping. In the rest condition, patients adopted a seated posture with their eyes open for 2 minutes. In the standing condition, patients alternated between 1 minute of upright posture on the force plates and 1 minute of seated posture, repeated 5 times for a total of 5 minutes standing. Similarly, in the stepping condition, patients alternated between stepping in place (while on the force plates) for 1 minute and seated posture for 1 minute, also repeated 5 times for a total of 5 minutes stepping. A minute typically refers to an artefact-minimal (see **Section 2.6.5**) period of 1 minute, that was confirmed visually, to mitigate the effect of movement artefacts.

The paradigm was performed twice: once in the OFF medication state and once in the ON medication state. The OFF condition was completed in the morning, with patients’ normal doses of levodopa withheld overnight. The ON condition was completed in the afternoon, with patients’ normal doses of levodopa administered at midday. This enabled a direct comparison between the two states.

### 2.4 Recordings

The surgical target was the STN. DBS systems from two companies were implanted: Medtronic Inc. Neurological Division, USA (octopolar directional leads, SenSight^TM^ model 33005) or Boston Scientific, USA (octopolar directional leads, Vercise^TM^ model DB-2202). Electrodes were implanted as previously described (Wiest et al., 2023), connected to temporary lead extensions and externalised through the temporal or frontal scalp. LFPs from the STN were recorded throughout the paradigm using the TMSi-SAGA amplifier (provided by TMSi, Netherlands), at a sampling frequency of 4096Hz. EMGs were simultaneously recorded using the same amplifier with the electrodes placed on four separate locations in bipolar configuration: *left gastrocnemius (GA_L_)*, *left peroneus longus (PL_L_)*, *right gastrocnemius (GA_R_)*, *right peroneus longus (PL_R_)*. Peroneus longus was selected as a target area due to its key role in foot and ankle stability, which is critical for propulsion, balance, and postural control. Gastrocnemius was selected due to its significant role in gait, such as influence on speed and power, propulsion, and control of important joints. Additionally, two force plates were recorded using the same system in order to capture the phase of the steps. A total of six EEG’s were also placed on the sites: C_z_, C_3_, C_4_, C_Pz_, C_P3_, C_P4_, to capture cortical activity. The ground electrode was placed on the participants wrist.

### 2.5 Contact selection

Bipolar configuration was applied to LFP recordings in post-analysis to reduce activities from volume conduction and to focus on locally generated activities. Several different bipolar configurations were created post-hoc based on unipolar LFP recordings from neighbouring contacts. From these, continuous wavelet transform (CWT) was utilised to determine the amplitude spectral density (ASD) of all bipolar signals from the OFF medication rest condition. The bipolar configuration with the largest beta activity was selected as the configuration for use in the analysis.

### 2.6 Data processing

The following analysis pipeline was implemented primarily in MATLAB (version 2019b). SPSS was used for ANOVA, and Spike2 for visualisation.

#### 2.6.1 Stepping analysis

Stepping frequency was defined as the number of steps completed per second. This was calculated by counting the number of steps (by observing the number of times the amplitude of the force plate exceeded a threshold) during the artefact-free part of the trial (see **Section 2.6.6**) and dividing this value by the number of seconds. This was computed for each foot and an average stepping frequency was obtained for each participant. The stepping frequency variance of each participant was defined as the variability of the stepping frequency over the trial. This was calculated by splitting the condition into 10 s segments and calculating a stepping frequency for each segment. The standard deviation of these stepping frequencies over all segments for each patient and each condition was then used to quantify variability. For analyses focusing on the stepping frequency variability during the first and last parts of the condition, the stepping duration of each step was recorded, and the standard deviation was calculated separately for the first 20 s and the last 20 s.

#### 2.6.2 LFP and EMG analysis

Continuous wavelet transform (CWT) was used for time-frequency decomposition of the chosen bipolar LFPs in the STN and the EMG activities from the recorded muscles. The data was pre-processed with a 100 Hz low pass filter, a 1 Hz high pass filter (both second order, two-pass Butterworth filters), and a 50 Hz notch filter to eliminate line noise. CWT was used for time-frequency decomposition with a Morlet wavelet of 10 cycles and a standard deviation of 3. The amplitude of each frequency band at different time points was calculated by taking the absolute value of the complex output. The average amplitude for different frequency bands and task conditions were then quantified. The amplitude of each individual frequency (per 1 Hz) was z-scored over each condition (ON and OFF) for each participant and then mean averaged over the period and frequency band in question.

#### 2.6.3 STN beta burst analysis

Beta band activity was defined within the frequency range of 12–35 Hz (Jurewicz et al., 2018; Moran et al., 2011). Beta bursts were subsequently defined as time periods where the average beta amplitude exceeded its 75th percentile for a minimum of 200 ms (Tinkhauser et al., 2017, 2018). This 75th percentile was calculated for each condition separately, meaning there were differing raw thresholds for bursts.

#### 2.6.4 IMC and STN-muscle coherence

The phase–locking value (Aydore et al., 2013; Celka, 2007) was used to calculate STN-muscle coherence and intermuscular coherence. This was to compute the phase consistency between the STN and the lower extremity muscles, as well as the IMC between these muscles. The PLV provides estimates of synchrony independent of the amplitude of oscillations. This is in contrast to measures of coherence where phase and amplitude are intertwined (Uhlhaas et al., 2010). To calculate PLVs, the signals of interest were first band-pass filtered using a digital IIR filter, prior to Hilbert transformation. The instantaneous phase of each signal at each time point was extracted, and the phase difference between the signals were calculated. The vector strength of the phase difference was computed using a sliding window technique with a fixed window length of 250 ms period, with 125 ms before the sample and 125 ms after. The value at each time point is the vector strength of the phase difference over this 250 ms window. This was computed over the entire duration of the condition under analysis. This procedure was repeated for each frequency band to generate a time-frequency coherence plot. The mean was found for each participant, before finally computing the mean across all participants.

For STN-muscle coherence, the STN signal was kept the same and the EMG data was randomly shuffled to generate a comparison which was subtracted from the original to give the difference between the observed data and the shuffled data. This eliminates the influence of one signal and focuses on the coherence between the two of them.

#### 2.6.5 Stepping phase related modulations

To analyse the stepping phase related changes in the STN LFPs, EMG activities, as well as the STN-muscle connectivity and IMC, the time-series of the recorded stepping force of each foot was first Hilbert transformed to find the phase of the step, from -π/2 to π/2 radians. Each step cycle was then divided into 181 different bins according to the calculated phase. The average amplitude of STN LFP activities, muscle EMG activities, STN-muscle connectivity, and IMC in different frequency band were found for each phase bin. These were z-scored for each participant, by condition (ON and OFF), as described in **Section 2.6.2**. This result was then averaged over all participants, allowing for analysis of electrophysiological modulation according to stepping phase.

#### 2.6.6 Trial Rejection

Prior to processing in MATLAB, the raw recorded data was loaded into Spike2 for visualisation. Obvious artefacts were identified and only data considered as clean was selected for inclusion in the analysis. Artefacts include ocular, jaw clenching, or mechanically induced (from the cable movement) disturbances on the salient channels for the analysis, particularly the STN LFPs and EMG data. In addition, clearly identifiable stepping induced force readings from the plates is also required to define data from that step as clean.

#### 2.6.7 Statistics

There were two main statistical tests utilised in the study. The first was the repeated measures ANOVA, which was adopted when the effects of more than two experimental conditions (for example medication, frequency band, locomotion status) were tested. The second was the Wilcoxon Signed Rank test for when there were two groups, and for post-hoc testing. As it is a non-parametric test it is more robust to the small sample sizes in this work.

Permutation-based cluster analysis was employed to test whether the EMG amplitude, STN-muscle coherence, and IMC in different frequency bands during stepping were significantly different between ON and OFF levodopa medication, or between beta bursts and no bursts. This was implemented by generating a paired *t*-test plot between the two conditions. *T*-statistics for the clusters were then determined, as well as the sizes of the clusters in pixels, with the largest values selected as the largest cluster. Then, the null distribution was generated by randomly swapping the pairings in the paired samples *t*-test, where the largest *t*-statistic and cluster size were recorded for each permutation, which created a set of possible cluster sizes under the null hypothesis. Finally, two p-values were generated, one for the cluster size and one for the accumulated *t*-statistic, computed by comparison with the null distribution. Only the overall *p*-value from the accumulated *t*-statistic method is reported here, because there were no results that changed between the two methods.

## 3. Results

### 3.1 Levodopa decreases variance of patient stepping frequency

To evaluate behavioural changes induced by the medication, analyses were conducted on patient stepping patterns (i.e. stepping frequency and its variability). Results demonstrate that there was a borderline significant effect of medication on the stepping frequency, with a marginal increase in the frequency when ON medication (*Z = −1.73, p = .084*; **Fig. 1B**). Furthermore, an analysis of the variability (standard deviation) of stepping frequency revealed a statistically significant difference between conditions (*Z = 2.29, p = .022;* **Fig. 1A**), demonstrating reduced variability when ON medication.

**Figure 1:**
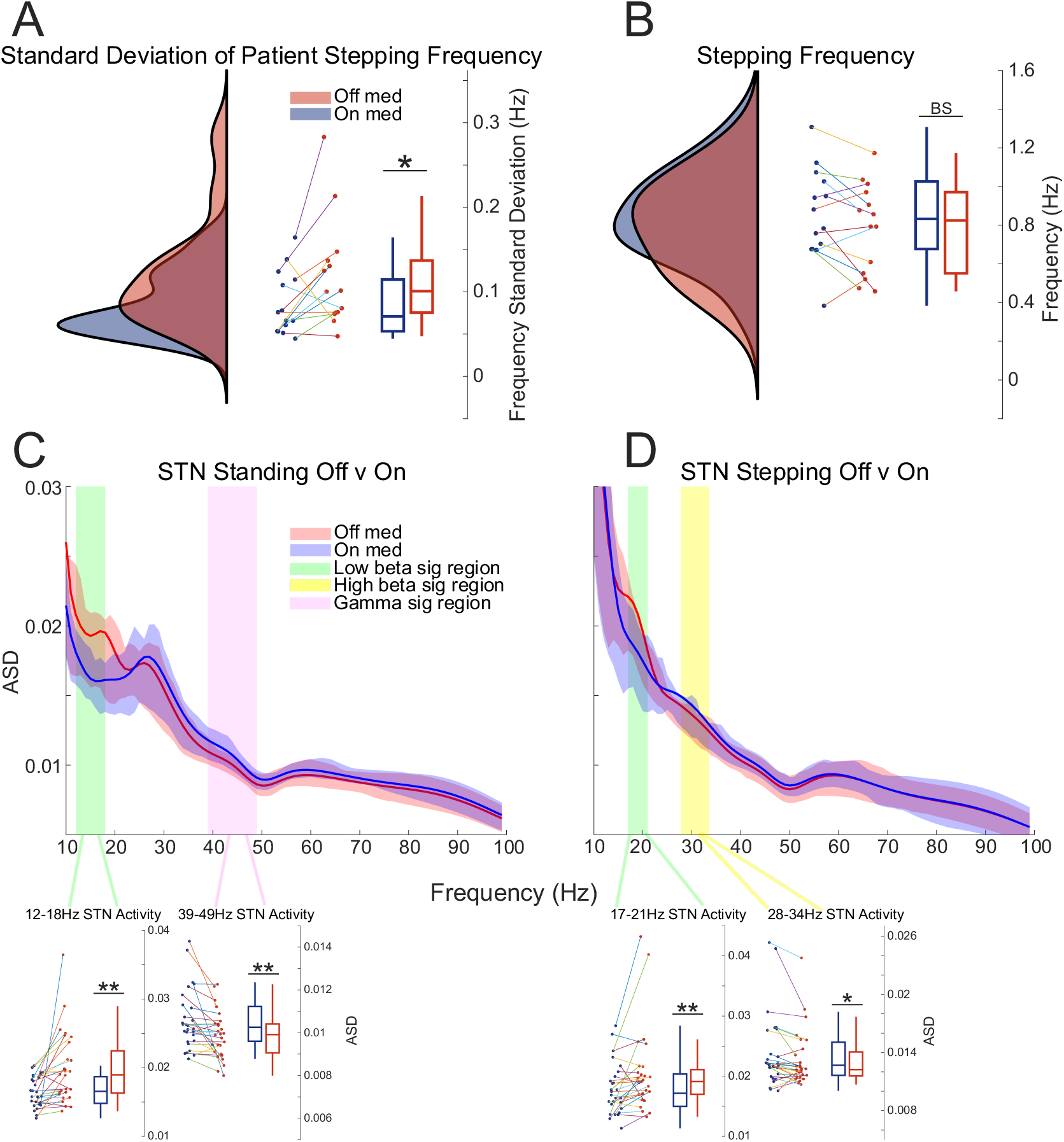
Overview of the experiment. A boxplot of the standard deviation of the patients’ stepping frequency is presented in **A** while the patients’ stepping frequency is depicted in **B**. The amplitude spectral density of activity in the STN over the standing and stepping trials is presented in **C** and **D** respectively.

A two-way ANOVA was conducted to evaluate how medication (ON and OFF) and time-related changes affect stepping frequency and its variability, which was studied by examining specific time intervals (initial 20 s and final 20 s of stepping). This tests whether the overall increase in the variability of the step frequency when OFF medication was due to changes over time. This analysis shows that medication had a marginally significant effect on the stepping frequency (*F(1, 13) = 4.21, p = .06*), increasing it on average. It also shows that there was a statistically significant effect of the time interval in the task (*F(1, 13) = 8.273, p = .013*), where stepping frequency had increased in the final 20 s. In addition, no interaction effects were observed between the medication and time interval (*F(1, 13) = 0.005, p=.94*), suggesting that the stepping frequency increases with time in the task independent from medication. For variability, there was no effect of medication on the standard deviation of stepping frequency in the first and last 20 s of stepping (*F(1, 13) = 3.306, p = .092*). The time interval did not affect the stepping frequency standard deviation (*F(1, 13) = 0.030, p = .865*) and no interaction effects were observed with medication (*F(1, 13) = 0.181, p = .678*). This implies that the variability observed in the stepping frequency was not due to time-related changes (slowing down or increasing) in the step frequency within each recording block.

3.2 Different effects of levodopa and locomotor status on low- vs. high-beta band activities in the STN LFPs

Initial analysis examined the effects of medication on STN LFPs during standing and stepping using a permutation cluster analysis, as presented in **Fig. 1C and D**. The amplitude spectral density was computed for frequencies ranging from 10 – 100 Hz and normalised by dividing each value by the area under the curve. During stepping, this analysis revealed a significant reduction in the 17-21 Hz band when OFF medication (*t = 2.66, p = .011*), as well as an increase in 28-34 Hz activity (*t = −2.30, p = .017*). During standing, activity in the 12-18 Hz band was significantly decreased (*t = 3.92, p < .001*) while activity in 39-49 Hz slightly increased (*t = −3.46, p < .001*). Subsequently, a three-way ANOVA was conducted on the medication condition (ON vs. OFF), the locomotion status (stepping vs. standing), and the beta frequency band *(high (21-35 Hz)* vs. *low (12-20 Hz)),* to analyse the effect on the beta power. This analysis reveals that there was a significant effect of the medication condition (*F(1, 13) = 8.40, p = .007*), the locomotion status (*F(1, 13) = 8.72, p = .006*), and the beta band (*F(1, 13) = 32.38, p < .001*), on the beta amplitude. Interaction effects demonstrate that medication and locomotion status do not interact with each other (*F(1, 13) = 0.78, p = .39*). However, some interaction effects were recorded with medication influencing the effect of beta band on the beta amplitude (*F(1, 13) = 8.87, p = .006*), and locomotion status influencing the effect of beta band on beta amplitude (*F(1, 13) = 33.38, p < .001*). There were no interaction effects between the medication, motor, and beta band conditions (*F(1, 13) = 0.092, p = .346*).

Post-hoc analyses revealed that there was a significant effect of levodopa on the high beta band amplitude during standing (*Z = −2.08, p = .037*) and stepping (*Z = −2.05, p = .040*), indicating that levodopa increases high beta oscillations. During standing, there was also a significant effect of levodopa on low beta band amplitude (*Z = 3.61, p < .001*), but the direction of this change was opposite to that observed in the high beta frequency band, with levodopa reducing low beta activity. Notably, this effect was not observed during stepping (*Z = 1.622, p = .105*).

Additional post-hoc analyses showed that high-beta amplitude was significantly lower during stepping than standing, regardless of medication state (OFF medication: *Z = 3.73, p < .001*; ON medication: *Z = 4.06, p < .001*). On the contrary, low-beta band amplitudes were higher during stepping compared to standing, both when OFF medication (*Z = −3.33, p < .001*) and ON medication (*Z = −3.53, p < .001*). These different effects of locomotor status on low- vs. high-beta activity suggest distinct pathophysiological roles of these frequency bands in Parkinsonian stepping.

### 3.3 Decreased beta band activities in the STN LFPs and EMGs around the contralateral late-stance and pushing-off phase when ON levodopa

To explore the effect of stepping phase on STN and EMG activity, and whether medication induced phase-specific changes, force plate readings were used to extract the step phase, as detailed in **Section 2.6.1 (Fig. 2A**). Consistent with findings from previous studies, step-phase related modulation of beta band activities in the STN LFPs were observed. These modulations mirrored contralateral muscle activity both ON and OFF medication (**Fig. 2B-K**), with beta band activities in the STN LFPs increasing and decreasing in sync with muscle activities in the contralateral leg. One-dimensional permutation cluster analysis demonstrated a significant difference in STN beta activity (12-35 Hz) from 0 to π/4 between OFF and ON medication states (*t = 2.299, p = .018*, **Fig. 2D**). This phase corresponds to weight-shifting and movement initiation (pushing-off) stages of the gait cycle, where beta activity is generally reduced. This beta reduction was more pronounced in the ON medication state, indicating a stronger suppression of pathological beta activity during the contralateral late-stance and ‘pushing-off’ phase compared to the OFF medication state. The same analysis was then conducted for the EMG activity, which revealed a clear increase in muscle activity during the same 0 to π/4 phase in both muscles (*gastrocnemius: p = .030, peroneus longus: .043*) (**Fig. 2H, I, L, and M**). While this increase of activity was observed across beta and into the gamma range, the exaggerated beta activity is consistent across muscles and particularly prominent in the gastrocnemius (**Fig. 2J, K, N, O**).

**Figure 2:**
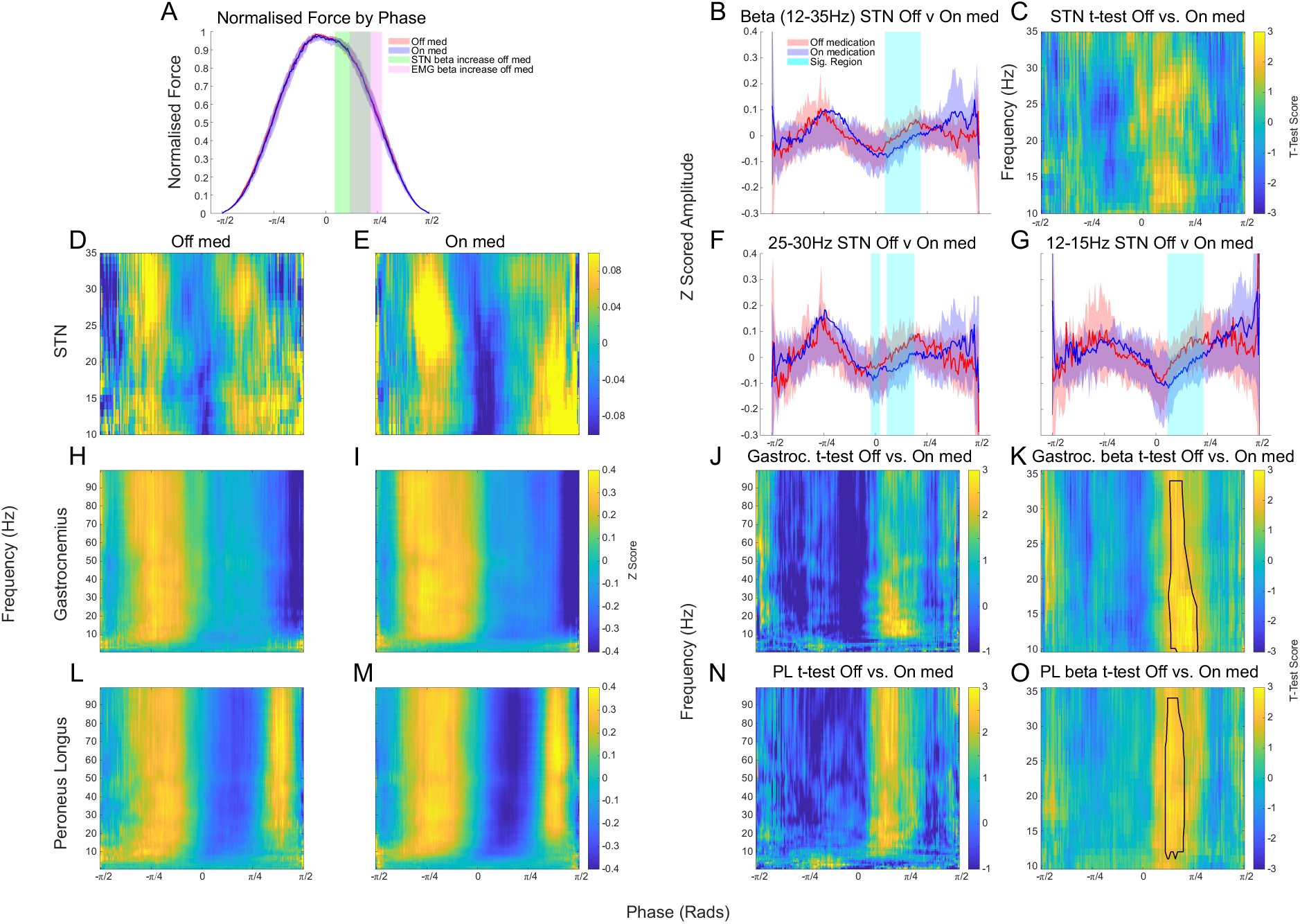
Analysis of electrophysiology by stepping phase. The normalised force of the plates by the stepping phase, computed using the Hilbert transform, is depicted in **A**. The time frequency decompositions of the STN, gastrocnemius, and peroneus longus OFF medication are displayed in **B**, **F**, and **J**, while ON medication items are displayed in **C**, **G**, and **K** respectively. Data are normalised within condition, i.e. separate normalisation for each condition. **D** depicts the beta activity averaged over 12-35 Hz for OFF and ON medication, while the *t*-test scores between OFF and ON in the time frequency domain in the STN, gastrocnemius, and peroneus longus are shown in **E**, **I**, and **M** respectively. The *t*-test scores between OFF and ON from 1-100 Hz in the gastrocnemius are presented in **H** for the gastrocnemius, and **L** for the peroneus longus.

### 3.4 Increased beta-band STN-muscle and intermuscular coherence during late stance phase, with the IMC further increased by levodopa in the same time window

Further analysis examined STN-muscle coherence and intermuscular coherence (IMC between the gastrocnemius-peroneus longus coherence of the same leg) in relation to the stepping phase. The step-phase related modulation pattern of the IMC (**Fig. 3A-C**) showed an opposite pattern compared to the amplitude modulation pattern (**Fig. 2J&N**). The STN-muscle coherence (average of STN-gastrocnemius with STN-peroneus longus) and IMC in the alpha/beta bands increased from phase 0 to π/4 (i.e. the late-stance and pushing-off phase), while amplitudes of STN and muscle activities decreased relative to phases in the step cycle. Permutation cluster analysis revealed a significant increase in IMC in the beta band (*p = .020*) between 0 and π/4 when ON medication compared to OFF (**Fig. 3A**). This indicates that levodopa enhances phase synchrony between the gastrocnemius and peroneus longus muscles in the same leg during late-stance and pushing off phase, despite the overall reduction in muscle activity observed in the ON medication state.

**Figure 3:**
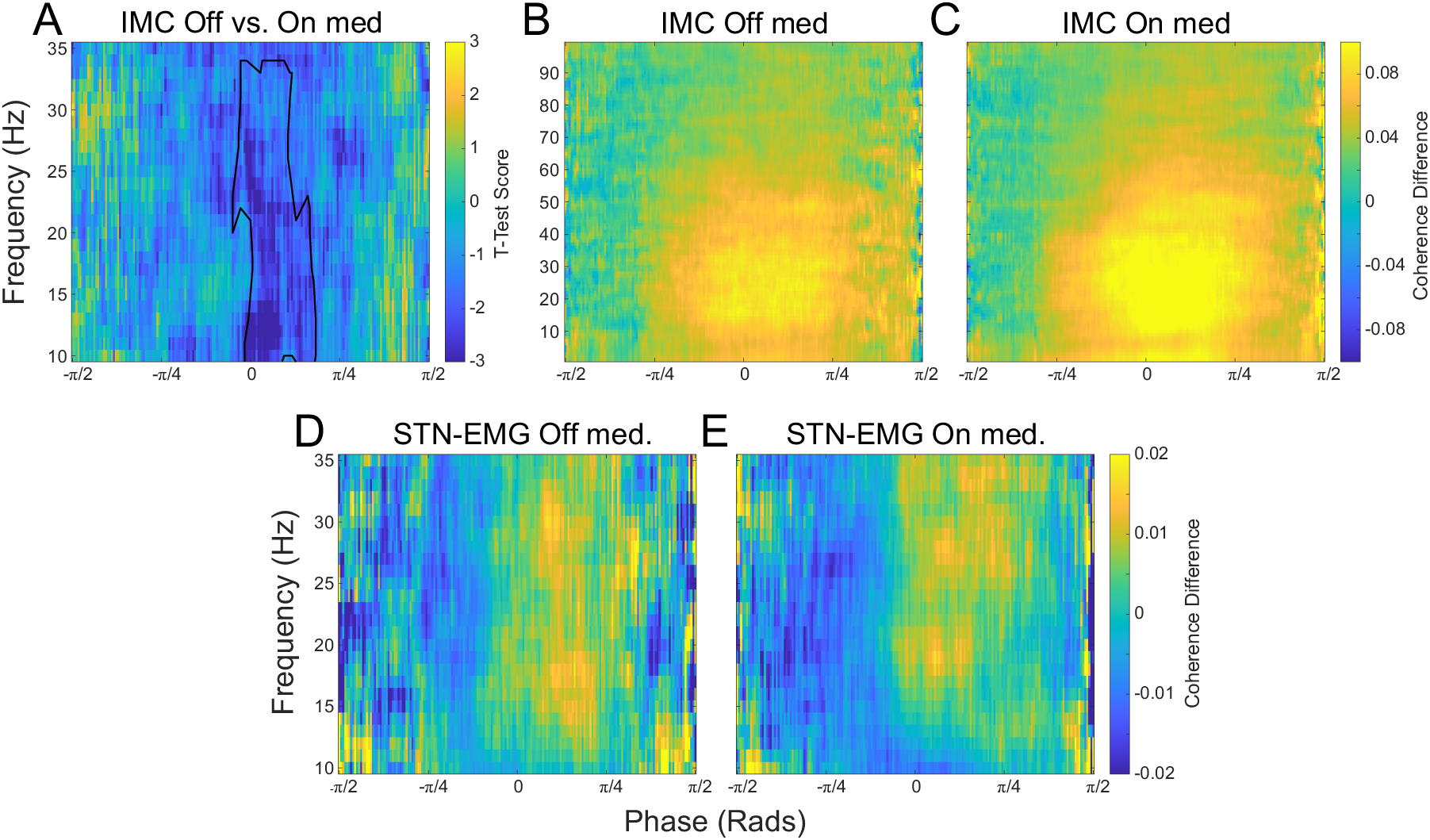
Intermuscular coherence (IMC) by stepping phase between the gastrocnemius and peroneus longus muscles of the same leg presented in **A**, **B**, and **C**, and STN-EMG coherence by stepping phase medication presented in **D** and **E** respectively. Time frequency maps presented for OFF medication in **B** and **D** and ON medication in **C** and **E**, and differences between OFF and ON medication for IMC with the *t*-test score in **A**. For **B**, **C**, **D**, and **E**, the coherence difference is computed by first calculating the phase locked coherence, and then subtracting a permuted version where the phase locked coherence is recalculated but with permuted EMG signal (for IMC only one of the signals is permuted).

### 3.5 STN beta bursts are linked to increased beta band activities in the EMG

Beta bursting activity was extracted according to the procedure outlined in **Section 2.6.3**. In addition, the amplitude by frequency of the EMG was computed over the entire recording. Then, EMG activity corresponding to STN beta burst onset was extracted. Due to the previous finding showing phase-locked increases in EMG activity during a specific phase of the stepping cycle, only STN bursts occurring between 0 and π/4 radians were considered. A permutation cluster analysis revealed a significant increase in EMG beta band amplitudes around STN beta burst onset between OFF and ON medication for both the gastrocnemius (*p = .020*; **Fig. 4C**) and the peroneus longus (*p = .040*; **Fig. 4D**).

**Figure 4:**
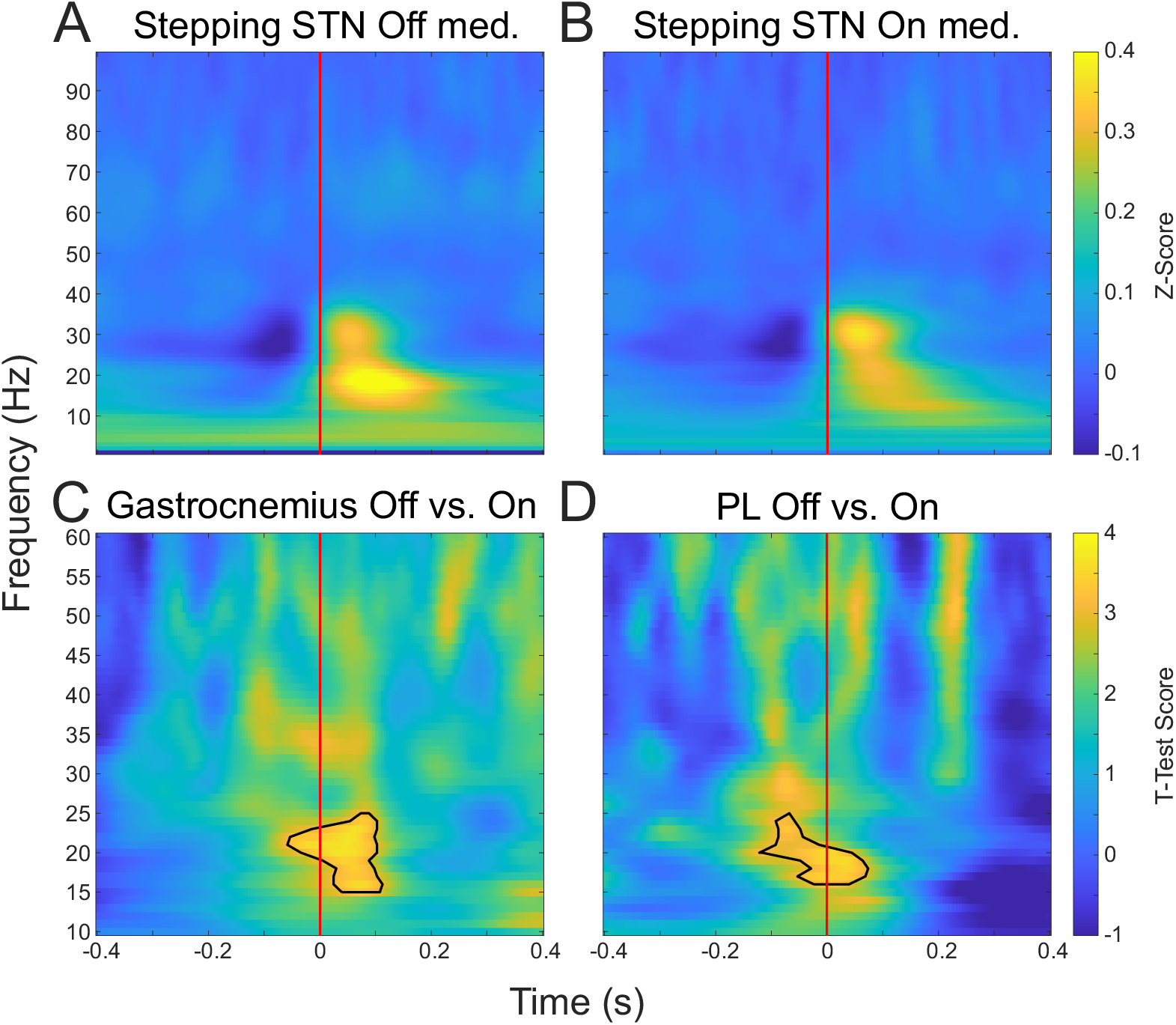
Effect of STN beta bursts from phases 0 to π/4. Beta bursting activity is extracted where the amplitude exceeds the 75^th^ percentile (indicated by the vertical red line in the figures), shown for OFF medication in **A** and ON medication in **B**. The concomitant activity is selected for the gastrocnemius and peroneus longus, with the *t*-test score between OFF medication and ON medication computed and presented in **C** and **D**.

## 4. Discussion

This study provides key insights into the role of beta oscillations during stepping in PD and the modulatory effects of levodopa. Firstly, levodopa medication and locomotion status (stepping vs. standing) were found to have opposing effects on low- vs. high-beta activity, indicating different roles of these beta sub-bands in the pathophysiology of gait in PD. Secondly, a step-phase specific effect of levodopa on beta oscillations during stepping was observed, with significantly decreased beta activity in the STN, the gastrocnemius, and the peroneus longus muscles during the late-stance and pushing off phase when ON levodopa medication. Lastly, STN beta bursts during this stepping phase were associated with increased beta activity in the EMGs in the gastrocnemius and peroneus longus muscles. Increased beta activity in the STN and lower extremity muscles may be associated with reduced movement efficiency, increased exertion, and impaired motor coordination. Together, these findings emphasize the importance of phase-specific beta modulation and its relationship with muscle activation, providing a potential rationale for targeted phase-based stimulation strategies to address gait impairments in PD.

### 4.1 Different effects of locomotion status (stepping vs. standing) on average amplitude of low- vs. high-beta frequency bands

The results demonstrate opposing effects of locomotion status (stepping vs. standing) on the average amplitude of low- vs. high-beta frequency bands in the STN. Specifically, stepping was associated with increased low-beta and reduced high-beta activity compared with standing, independent from the medication status. High-beta activity may be important for static posture control during standing, but its suppression might be necessary during stepping to facilitate active alternating movements, such as lifting, swinging, and pushing-off. This is consistent with a previous study showing that lower limb movements were associated with greater desynchronization in the high-beta frequency bands (24–31 Hz) compared to low-beta band or upper limb movements (Tinkhauser et al., 2019). Meanwhile, an earlier study has shown that PD patients with FOG exhibited elevated high-beta power than those without FOG, with significant reductions in high-beta power following levodopa administration along with suppression of FOG (Toledo et al., 2014).

The present findings also reveal different effects of levodopa on low- vs. high-beta frequency bands in the STN. Levodopa reduced low-beta oscillations but increased high-beta oscillations. The reduction in low-beta activity is consistent with previous findings (Giannicola et al., 2010; Jenkinson & Brown, 2011; Little & Brown, 2014), highlighting its pathological nature. On the other hand, the increase in high-beta activity suggests a potential physiological function of this frequency band during standing and stepping. Recent research supports the notion that supplementary motor area activity selectively drives high-beta STN activity via the hyperdirect pathway, suggesting a functional role for high-beta frequencies in cortico-subcortical communication (Oswal et al., 2021). Furthermore, low- vs. high-beta band cortico-subcortical coherence have been implicated to play different roles in movement inhibition and expectation (Cao et al., 2024).

It should be recognised that most patients recorded in this study are tremor dominant, or have main symptoms of bradykinesia and rigidity, therefore, it was not possible to conduct analyses of patients with FOG vs. those without FOG. Further studies focusing on patients with FOG may further elucidate the pathophysiological role of low- vs. high-beta in gait impairment in PD.

### 4.2 Step-phase specific effect of levodopa on STN LFPs, EMGs and intermuscular coherence during stepping

Consistent with previous findings, there was step-phase related modulation in beta band activity in the STN, which increased during the early stance phase and decreased during the late-stance and when pushing off from the contralateral leg. This reduction in beta activity during the late-stance and pushing off phase may be linked to the movement initiation (Crenna et al., 2006; Fischer et al., 2018; Kojovic et al., 2016). Yet, the role of muscle activity during stepping is less understood. In the presented study, intermuscular coherence increased during the late-stance and pushing-off phase, despite a reduction in total amplitude of muscle activity, which may indicate more effective coordination between the recorded muscles during movement initiation.

In addition, a step-phase specific effect of levodopa on STN LFPs, EMGs and intermuscular coherence during stepping was observed, which was constrained to the late-stance and pushing-off phase. When OFF medication, there was an increase of STN beta activity and EMG activities, as well as reduced intermuscular coherence in the effected phase. Increased STN beta activity in this phase window may be associated with slowing down in movement initialisation and reduced movement speed (He et al., 2020; Lofredi et al., 2019; Torrecillos et al., 2018). On the other hand, increased EMG activities in both the gastrocnemius and peroneus longus were observed, but with a reduction of intermuscular coherence (phase synchrony) in the beta frequency band. The reduction in STN beta activity appears to precede changes in EMG activity (**Figure 2A**), suggesting a causal relationship where STN activity influences muscle activation. This is further supported by the observed effect of beta bursts on EMG activity, which revealed an increase in muscle activation following heightened beta activity in the STN, similar to previous observations of cortical beta bursts (Echeverria-Altuna et al., 2022; Simpson et al., 2024). Conversely, reduction of STN beta activity could enable the muscle to release from a tonic state and transition more smoothly into the step phase.

Although the increase in muscle activity resulting from STN beta bursts may initially seem minor, it may have significant consequences for gait. One possible outcome is a reduction of movement efficiency, requiring greater effort to complete the same movement (Graham et al., 2015), which could result in faster depletion of energy levels (Hagell & Brundin, 2009), further slowing movement and impairing motor coordination. Increased muscle activity may also inhibit natural stepping motion, which may result in compensatory mechanisms and strategies that are suboptimal (Nonnekes et al., 2019), such as recruitment of other muscle groups which could also reduce energy levels (Fletcher & MacIntosh, 2017). Furthermore, this increase of muscle activity can cause co-activation of multiple muscles and altered muscle activation patterns, which are both linked to altered timing of the gait cycle (Srivastava et al., 2019). In turn, this alteration of timing has been proposed as a possible contributor to FOG (Nieuwboer et al., 2004). Similarly, the inability of patients to effectively produce muscle coordination at movement initiation may also contribute to stepping disturbance. This could result in a number of consequences including an increase in the effort to execute the movement, prompting of compensatory mechanisms, and increased levels of fatigue.

### 4.3 Implications for future stimulation strategies

There are several implications for future stimulation strategies that emanate from this work. The results suggest that during parkinsonian stepping the motor outputs (both beta band activities in the STN LFPs and EMG activities in the lower extremity) were significantly affected from phases 0 to π/4 in each step cycle, which corresponding to the late-stance and pushing-off phase. The results support the view that stimulating between 0 and π/4, utilising a stepping-based phase-triggered stimulation strategy, could further probe the causal relationship between the observed increase in the beta band activities in STN LFPs in this time window during OFF medication condition and gait impairment in PD. It may also offer some beneficial effects for patients during stepping and potentially free walking. It could also reduce variations of timing in the gait cycle, alleviating one of the contributing factors to freezing of gait, potentially reducing these episodes. Another possible ramification is a reduction in reliance on compensatory mechanisms.

### 4.4 Limitations and future work

One limitation of the work is that stepping does not necessarily translate directly to free walking, although some studies suggest STN activity is similarly modulated during free walking (Fischer et al., 2018). In addition, due to the challenge of recruiting patients with externalised leads, the sample size is limited and most patient have main symptoms of tremor, bradykinesia and rigidity. Therefore, the effect of levodopa on stepping and associated changes in electrophysiology was the focus, and it was not possible to study FOG specifically. Another limitation is the timing of the study, as the OFF medication sessions are recorded in the morning and the ON medication sessions in the afternoon, patients may have had less energy and motivation for the ON medication sessions.

Future work could focus on STN activity around periods that can instigate freezing episodes for example, stopping, turning, or moving through a doorway. Furthermore, works that test different stimulation strategies at different phases of stepping are also critical to evaluate the efficacy of the proposed approach, with testing in real-world gait scenarios a prerequisite for clinical translation.

## Funding

This work was supported by the Medical Research Council (MC_UU_0003/2, MR/V00655X/1, MR/P012272/1), the Medical and Life Sciences Translational Fund (MLSTF) from the University of Oxford, the National Institute for Health Research (NIHR) Oxford Biomedical Research Centre (BRC), Parkinson’s France, and the Rosetrees Trust, UK.

## Competing interests

The authors declare no competing interests.

---

**Table 1:** This table gives some information about the patients in the study. Firstly, patient ID is given, along with the type of DBS leads, and the surgical target. The patients’ dominant hand (DH) is also shown, as is their sex assigned at birth (M/F), age, disease duration (Dis. Dur), and predominant symptoms before surgery.

| Patient ID | DBS Leads | Target | DH | M/F | Age | Dis. Dur | Predominant Symptoms before surgery |
| --- | --- | --- | --- | --- | --- | --- | --- |
| P001 | Medtronic Sensight (directional) | STN | R | F | 63 | 9 | Akinetic-rigid |
| P002 | Medtronic Sensight (directional, Percept IPG) | STN | L | M | 75 | 6 | Tremor both hands, light rigidity |
| P003 | Medtronic Sensight (directional) | STN | R | M | 61 | 5 | Tremor, worst on right side |
| P004 | Medtronic Sensight (directional) | STN | - | M | 58 | 12 | Rigidity and bradykinesia |
| P005 | Boston Scientific (directional) | STN | R | M | 66 | 10 | Akinetic-rigid |
| P006 | Medtronic Sensight (directional) | STN | R | M | 57 | 5 | Tremor dominant worst on right, bradykinesia and FoG |
| P007 | Boston Scientific (directional) | STN | R | M | 63 | 13 | Akinetic-rigid |
| P008 | Boston Scientific Octode (non-directional, 2201 leads) | ViM/Vop + STN | R | M | 70 | 9 | Tremor dominant |
| P009 | Medtronic Sensight (directional, 33005 leads) | STN | R | F | 62 | 12 | Akinetic-rigid |
| P010 | Boston Scientific (directional) | STN | R | M | 69 | 30 | Tremor dominant |
| P011 | Medtronic Sensight (directional) | STN | R | F | 67 | 12 | Severe off symptoms with dyskinesia and rigidity, some right side tremor |
| P012 | Medtronic Sensight (directional) | STN | R | M | 67 | 6 | Tremor and rigidity in left side |
| P013 | Boston Scientific (directional) | STN | R | M | 61 | 14 | Tremor dominant |
| P014 | Medtronic Sensight (directional) | STN | - | F | - | - | Rigidity in left side |

## Notes

### Competing Interest Statement

The authors have declared no competing interest.

